# Quantifying variation by the ratio of CSS to UCSS

**DOI:** 10.1101/096784

**Authors:** M. Pahlevani

## Abstract

The variance, the average of squared deviations of data values from their mean, is the most widely used criterion for measuring the variation. Small amounts of the variance indicate that the values tend to be close to the mean and its high amounts are indicative of more dispersion around the mean. However, when dealing with a single variable or variables with different measuring units, variance can not give us a proper understanding of the actual extent of variation. RCU, the ratio of corrected sum of squares (CSS) to the uncorrected sum of squares (UCSS) can quantify variation too. The values of RCU vary from zero, for a situation in which all values are the same, to 100 % when they are completely different or symmetric. To compare the efficiency of RCU and variance for measuring variation, data of seven variables with different units of measurement for 17 wheat cultivars including yield (g/plant), spikes (numbers), height (cm), earliness (days), viability (%), hectoliter (kg/hectoliter) and EC (micromohs/cm) were used. Variance of these variables respectively was 83.34, 72.01, 353.23, 81.48, 5.31, 69.52 and 7167.47. The highest and lowest RCU was obtained for spikes (9.38 %) and viability (0.05 %), respectively. The RCU for yield, height, earliness, hectoliter and EC was 8.97, 2.67, 0.29, 1.30 and 2.85 %, respectively. The RCUs were somewhat similar to coefficient of variation of the variables. Based on the RCU, the extent of variation was medium for spikes and viability and was low for the other variables. As a result RCU could be used as a simple criterion to quantify the actual extent of variation in the data.

**Abbreviation:**

CSS: corrected sum of squares
UCSS: uncorrected sum of squares
RCU: the ratio of CSS to UCSS
CV: coefficient of variation
VAR: variance

## Introduction

Variation in plant genetic resources provides opportunity for breeders to develop improved new cultivars with desirable characteristics, which satisfy both researchers and farmers. Since the beginning of agriculture, human attention has been drawn to exploit the genetic diversity within crop species to meet subsistence food requirement, and now it is being focused to surplus food for growing populations. In biology, variation refers to every kind of observed differences among cells, living organisms or even among different species that might be due to genetic factors or because of the effect of environmental components on the expression of genetic potential. In the living world, variation is a tool help organisms for adaptation and survival against the environmental changes. The assessment of variation within and between plant populations is routinely performed using various techniques such as morphological, biochemical (allozyme), and DNA (or molecular). Variation (or dispersion, as it is sometimes called) is also the main requirement for selection superior types by natural selection as a basic element of the evolution. Variation into continuous and discontinuous is measured by different statistics such as quantiles, range, variance and standard deviation (Walpole et al., 2002). Due to some aspects, the variance and or standard deviation use in almost all studies related to measuring variation of the variables.

Variance, the second central moment, is the average squared differences of data from their mean. For a variable measured on the individuals σ^2^, variance calculates according to the below formula:

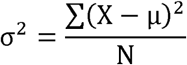

Although it is still unknown who proposed the formula for the first time, but term variance was first introduced by Ronald A. Fisher in 1918 (Fisher, 1918). The variance is the most widely used biometrical statistics in genetics and breeding of plants. The biometrical analysis such as ANOVA, regression, heritability, response to selection, gain from selection, selection index, combining ability, stability, clustering and factor analysis are directly calculated by the above formula of variance. According to the definition, the variance quantifies the amount of dispersion and shows how far the observations deviate from the mean (Zar, 2010). For a given population, variance of a variable is a non-negative numeric value that its amounts is an indicative of how the data points dispersed around the mean. However, if the variance is equal to zero indicates that all data values are identical.

The variance has square units. In most cases, the populations are so large that it is impossible to measure all of the individuals and the researchers attempt to estimate the variance by studying samples from the reference population. The variance of samples that draw out of the population is represented by S^2^ and the manner of calculation is similar to the variance with this exception that the numerator divided by N-1 instead of N, like the below formula (Jolicoeur, 1999):

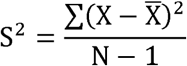

In practice, instead of using the above formula, below formula is used to calculate variance for samples:

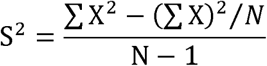

This formula is often referred to as a working formula or machine formula because of its computational advantages. Because of the widely application of the working formula; the components are named as below:

The numerator, ∑*X*^2^ – (∑X)^2^/*N*, is called corrected sum of squares or CSS in abbreviation. The CSS may also be visualized as a measure of the extent to which the data deviates from each other (Zar, 2010),

The subtractive component in the numerator, (∑X)^2^/*N*, is named correction factor or CF in abbreviation and the denominator, N-1, is called degree of freedom or simply DF. The numerator is again consisted of two parts which are, (∑X)^2^ the uncorrected sum of squares or UCSS, and CF. The CSS divides by DF to form variance or mean squares (MS). The term “mean squares” dates back at least to an 1875 publication of George Biddel Airy (Walker, 1929).

The most important feature of the variance is that it quantifies dispersion of the values around their mean. Of course, simplicity in calculation and its non-negative value are other valuable features of the variance as the most important and useful statistic for measuring the variation in all research fields. For example, consider the length of petiole in three plant samples is 0.1, 0.2 and 0.3 centimeters, thus UCSS, CF, CSS, DF and MS for this set would be 0.14, 0.12, 0.02, and 0.01 cm^2^, respectively. It means the values have dispersed with a 0.1 cm distance around the mean (*X*̅ = 0.2). The calculation was very simple since only three components, ∑ x, ∑x^2^, and N, are used in the working formula. As another hypothetical example, if the values of melting temperature for three metal alloys are 100, 200 and 300 °C, UCSS, CF, CSS, DF and MS would be 140000, 120000, 20000 and 10000 (°C)^2^, respectively. It shows, three metals have averagely dispersed with a 100 °C distance around the mean (*X*̅ = 200). In above examples, two variables with a quite similar dispersion come to mind because the relative differences of the numeric values in each variable are exactly the same. However, the calculations show that the level of dispersion quantified by the variance is very different for the two variables, 0.01 versus 10000. This indicates that the variance has some serious weaknesses. First of all, the variance of a variable has unit that is the square of measurement unit of the variable. For example, here its difficult to say whether 0.01 cm^2^ variation in petiole length of the plants is lower than 1000 (°C)^2^ variation in melting temperature of the alloys! Its usefulness is also limited because the unit is squared and not the same as the original measurements. The second weakness appears when only one variable is studying. In the above example, we do not know whether the variation of petiole length, which amount is 0.01 is really low? The same question arises for the melting temperature of the alloys, whether the variance of 10000 is really high? It has not been specified how much should be variance of a data set to be considered as high (or low). Actually, the numeric value of variance varying between zero and infinity does not provide information about the actual magnitude of variation of the data when considered by itself. It would be useful when compare to the variance of another sample for the same variable.

The first weakness could be solved by calculating the coefficient of variation (CV) which is a criterion without unit and declared in percent (Caulcutt, 1991). According to definition, CV is the ratio of the square root of the variance to the mean. The standard deviation is the positive square root of the variance; therefore, it has the same units as the original measurements. The term standard deviation was first coined by Karl Pearson in 1893, prior to which this quantity was called the mean error (Zar, 2010). The standard deviation of petiole length and melting temperature indicates the deviance of the data values from their mean is 0.1 cm and 100 °C, respectively. It is quite expected when differences of the data values with their mean is considered for each variable. CV of 50 % for petiole length and melting temperature was completely consistent with the equal extent of dispersion observed in these variables. The CV solves the unit weakness of the variance, however the second problem is still problematic and an absolute criterion is needed to measure the actual extent of dispersion.

The above examples clearly showed that quantification variation by the variance has serious weaknesses that cannot be solved by other criteria like standard deviation and CV. Therefore, developing a unitless measure that expresses the real extent of dispersion *per se*, while possess all features of the variance, especially the simplicity, is very necessary. It seems to quantify variation in a criterion without measurement unit, using the ratio of deviations of values from their mean is unavoidable.

Here it is assumed that RCU, the ratio of the corrected sum of squares (CSS) to the uncorrected sum of squares (UCSS), which is expressed as a percentage can be a suitable criterion for quantifying the amount of variation in samples and populations 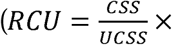 100). This paper will indicate the efficiency of RCU in comparison with the variance, especially for the aspects discussed above by using of data values for variables with different units of measurement. The main purpose of this study is whether RCU has the ability to quantify the variation in one variable as an absolute criterion or could be used as a relative criterion to compare the actual extent of variation for two or more variables. Also the effort will be made to investigate other features of this new measure, particularly in the aspects of simplicity in calculation and interpretation.

## Materials and methods

Seven variables, including grain yield per plant (g), number of spikes per plant, plant height (cm), earliness (days), grain viability (%), grain hectoliter (kg/hectoliter) and EC of grains (micromohs/cm) were measured on 17 wheat cultivars (Table 1). A three replicate experiment was done at the research farm and mean of the replications were applied in the data analysis. The study was performed at Gorgan University of Agricultural Sciences and Natural Resources, Iran, in 2014. Each experimental unit was comprised of six 120 cm lines which were planted 20 cm apart. Distance of seeds on the lines was 5 cm so that the final density was reached to 30 plants per square meter. The mean, variance (VAR), standard deviation, coefficient of variation (CV) and RCU of the measured variables was calculated. The following formula was used to calculate the RCU:

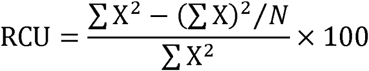

where, X and N are the variable (mean of three replicates) and number of the wheat cultivars (N=17), respectively. All calculations were performed with R software (R version 3.2.0) and Excel was used to produce graphs.

**Table 1:**
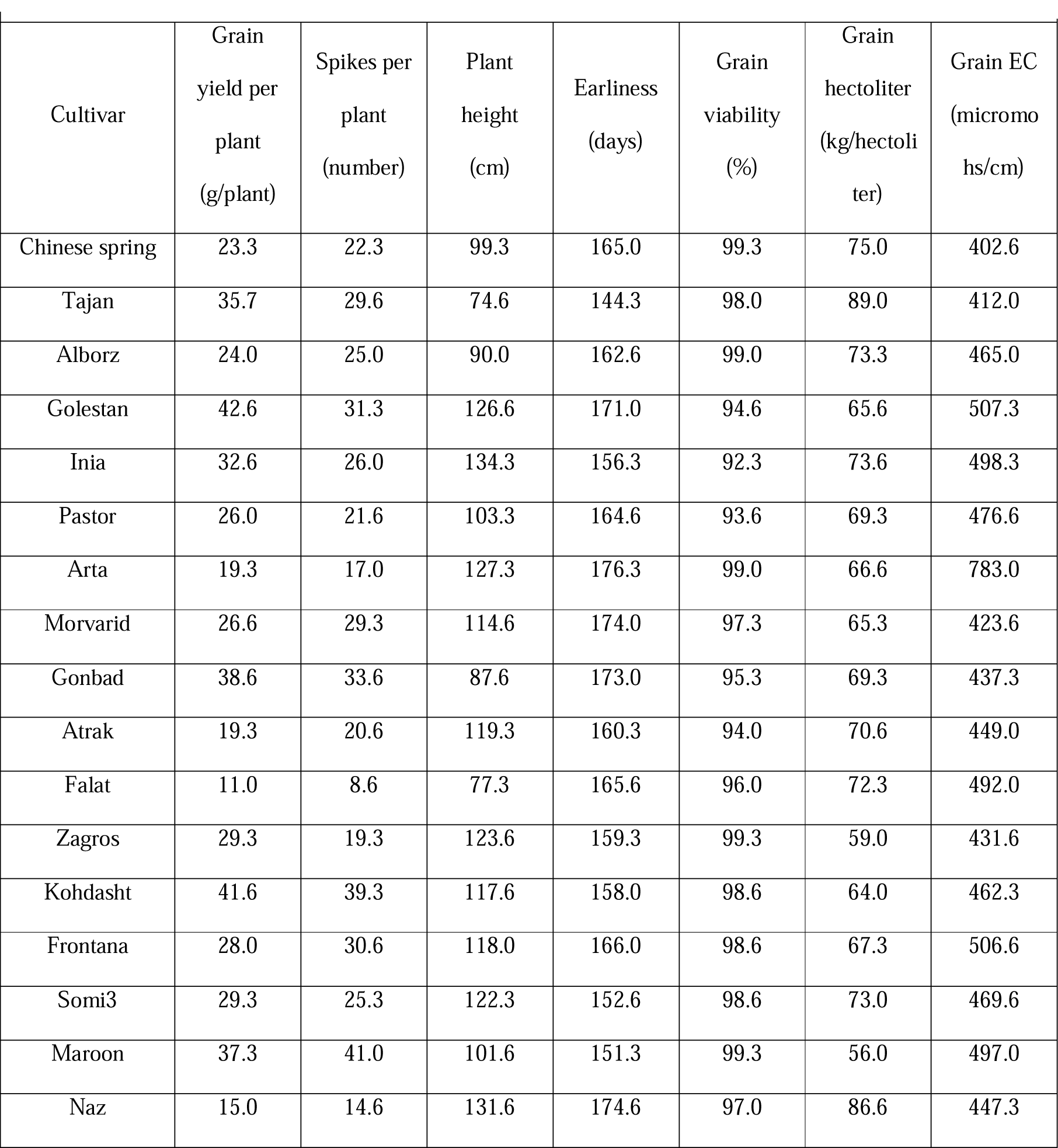
The mean of some morphological characteristics for 17 wheat cultivars.

|  |  |  |  |  |  |  |  |
| --- | --- | --- | --- | --- | --- | --- | --- |
| Cultivar | Grain<br>yield per<br>plant<br>(g/plant) | Spikes per<br>plant<br>(number) | Plant<br>height<br>(cm) | Earliness<br>(days) | Grain<br>viability<br>(%) | Grain<br>hectoliter<br>(kg/hectoli<br>ter) | Grain EC<br>(micromo<br>hs/cm) |
| Chinese spring | 23.3 | 22.3 | 99.3 | 165.0 | 99.3 | 75.0 | 402.6 |
| Tajan | 35.7 | 29.6 | 74.6 | 144.3 | 98.0 | 89.0 | 412.0 |
| Alborz | 24.0 | 25.0 | 90.0 | 162.6 | 99.0 | 73.3 | 465.0 |
| Golestan | 42.6 | 31.3 | 126.6 | 171.0 | 94.6 | 65.6 | 507.3 |
| Inia | 32.6 | 26.0 | 134.3 | 156.3 | 92.3 | 73.6 | 498.3 |
| Pastor | 26.0 | 21.6 | 103.3 | 164.6 | 93.6 | 69.3 | 476.6 |
| Arta | 19.3 | 17.0 | 127.3 | 176.3 | 99.0 | 66.6 | 783.0 |
| Morvarid | 26.6 | 29.3 | 114.6 | 174.0 | 97.3 | 65.3 | 423.6 |
| Gonbad | 38.6 | 33.6 | 87.6 | 173.0 | 95.3 | 69.3 | 437.3 |
| Atrak | 19.3 | 20.6 | 119.3 | 160.3 | 94.0 | 70.6 | 449.0 |
| Falat | 11.0 | 8.6 | 77.3 | 165.6 | 96.0 | 72.3 | 492.0 |
| Zagros | 29.3 | 19.3 | 123.6 | 159.3 | 99.3 | 59.0 | 431.6 |
| Kohdasht | 41.6 | 39.3 | 117.6 | 158.0 | 98.6 | 64.0 | 462.3 |
| Frontana | 28.0 | 30.6 | 118.0 | 166.0 | 98.6 | 67.3 | 506.6 |
| Somi3 | 29.3 | 25.3 | 122.3 | 152.6 | 98.6 | 73.0 | 469.6 |
| Maroon | 37.3 | 41.0 | 101.6 | 151.3 | 99.3 | 56.0 | 497.0 |
| Naz | 15.0 | 14.6 | 131.6 | 174.6 | 97.0 | 86.6 | 447.3 |

## Results and discussions

Mean of variables yield, spikes, height, earliness, viability, hectoliter and EC for seventeen wheat cultivars was presented in Table 1. It can be deduced from the means that, there were no similar cultivars for the studied variables (Table 1). Although there were more similarities for viability and earliness, for some variables such as yield and spikes the variability of the cultivars was most pronounced. The extent of variation for the studied variables was calculated using VAR, CV and RCU (Table 2). The largest VAR was observed in EC (7167.47 (micromohs/cm)^2^), and the lowest VAR was obtained in viability (5.31 %^2^). For other variables like yield, spikes, height, earliness and hectoliter, the VAR was 83.34 (square g/plant)^2^, 72.01 number^2^, 353.23 cm^2^, 81.48 days^2^ and 69.52 (kg/hectoliter)^2^, respectively (Table 2). Clearly understood that these variances do not meet the actual size of variation between cultivars (Table 1). The CV of yield, spikes, height, earliness, viability, hectoliter and EC was 31.40, 32.17, 5.36, 0.30, 11.50 and 17.11 %, respectively (Table 2). Detection of the highest CV for spikes and the lowest for yield shows that this criterion has well expressed the amount of variation in the measured variables. The highest RCU was observed in spikes (9.38 %) and the lowest RCU in viability (0.05 %) (Table 2). The RCU also for yield, height, hectoliter and EC was 8.97, 2.67, 0.29, 1.030 and 2.85 %, respectively (Table 2).

**Table 2.** Mean, variance, standard deviation, coefficient of variation and RCU for seven variables in wheat cultivars.

|  |  |  |  |  |  |
| --- | --- | --- | --- | --- | --- |
| Variable | Mean | Criterion of variation |  |  |  |
|  |  | variance | Standard<br>deviation | Coefficient of<br>variation (%) | RCU (%) |
| Grain yield per plant<br>(g/plant) | 28.21<br>(g/plant) | 83.34<br>(g/plant)<sup>2</sup> | 9.13<br>(g/plant) | 31.40 | 8.97 |
| Spikes per plant<br>(number) | 25.59<br>(number) | 72.01<br>(number)<sup>2</sup> | 8.48<br>(number) | 32.17 | 9.38 |
| Plant height<br>(cm) | 109.93<br>(cm) | 353.23<br>(cm)<sup>2</sup> | 18.79<br>(cm) | 16.58 | 2.67 |
| Earliness<br>(days) | 163.22<br>(days) | 81.48<br>(days)<sup>2</sup> | 9.03<br>(days) | 5.36 | 0.29 |
| Grain viability<br>(%) | 97.05<br>(%) | 5.31<br>(%)<sup>2</sup> | 2.30<br>(%) | 2.30 | 0.05 |
| Grain hectoliter<br>(kg/hectoliter) | 70.34<br>(kg/hectoliter) | 69.52<br>(kg/hectoliter)<sup>2</sup> | 8.34<br>(kg/hectoliter) | 11.50 | 1.30 |
| Grain EC<br>(micromohs/cm) | 480.07<br>(micromohs/cm) | 7167.47<br>(micromohs/cm)<sup>2</sup> | 84.66<br>(micromohs/cm) | 17.11 | 2.85 |

Depending on the magnitude of variation, the variance can include values from zero to positive infinity for types of variables. This issue is also apparent in Fig. 1 as the VAR for studied variables is much larger than the other two criteria i.e. RCU and CV (Fig. 1). Observation the largest VAR for EC (7167.47(micromohs/cm)^2^) was not also because of the high variation in this variable, since the highest variation was belonged to variable spikes as was shown by CV and RCU in Table 2. In addition, the variance of different variables could not be compared with each other, because they have their own measurement unit. For example, 7167.47 (micromohs/cm)^2^ VAR of EC can not be compared with 83.34 (days)^2^ VAR for earliness. This weakness is more pronounced for earliness and grain yield, two variables that had almost the same VAR (83.34 and 81.48, respectively), but with different unit of measurement. The weakness is well solved by the criterion CV. For example CV for variables yield and EC was 31.40 and 17.11 %, respectively, which obviously was in contrary to the VAR showing the variation in yield is more than EC. CV, coefficient of variation, proposed for the first time by Karl Pearson (1984) in order to standardize variation (David, 1995). The CV is the standard deviation of a variable that is expressed in terms of percentage of the mean. Because the standard deviation and the variable have identical units, CV has no units at all, a fact emphasizing that it is a relative measure, divorced from the actual magnitude or units of measurement of the data. The ratio is usually multiplied by 100% in order to express CV as a percentage. Based on the largeness of the CV, the studied variables ranked as spikes, yield, EC, height, hectoliter, earliness and viability (Table 2). However, the weakness is variable like spikes which had the highest CV, should not necessarily have the highest variance. By definition, CV could be a numeric value from negative infinity to positive infinity and in the cases that the mean is near to zero, the value of CV reacts largely in response to a tiny change in the variation and even sometimes reaches to infinity as well. Also, in cases where the mean is zero, CV is not calculable.

**Fig 1.**
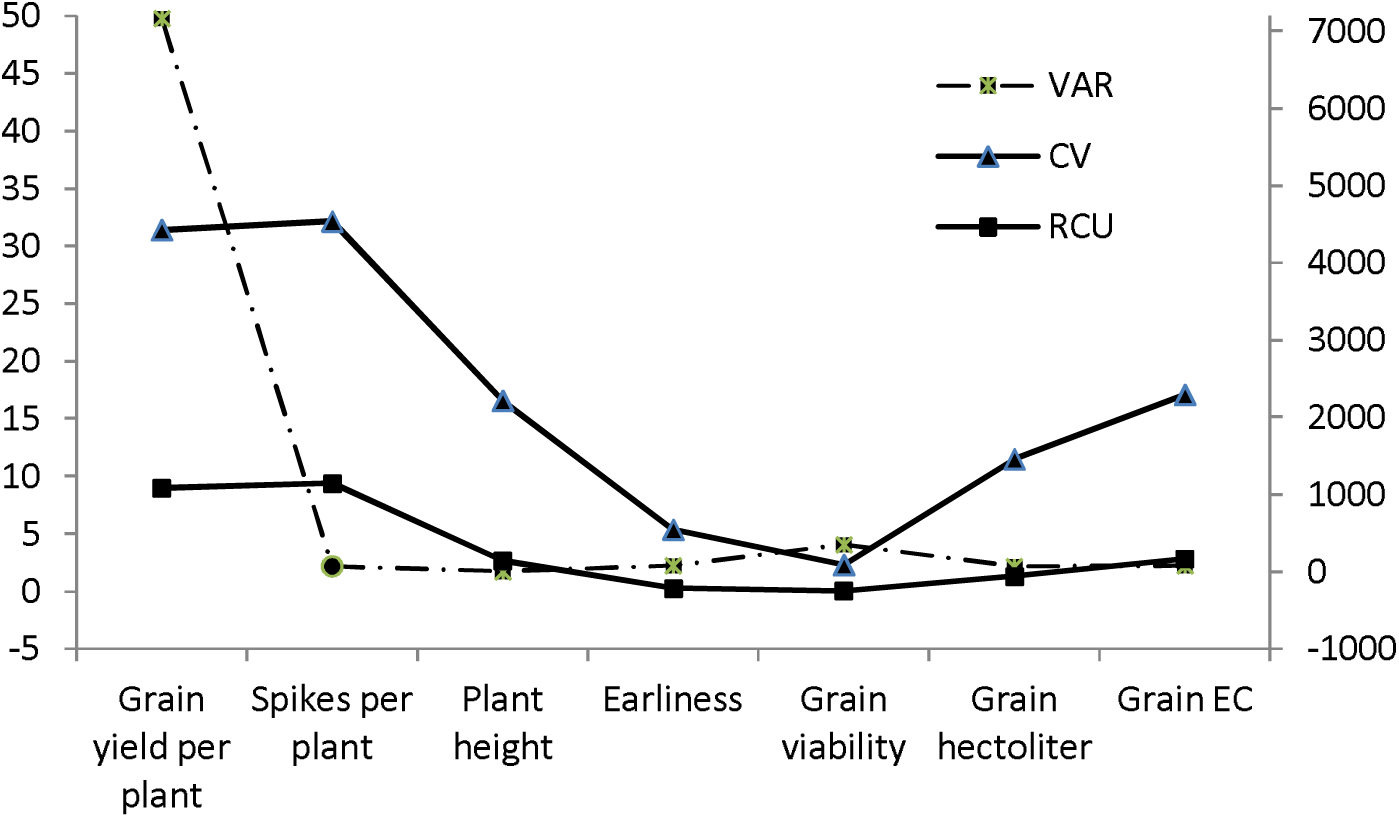
The correspondence of various dispersion criteria (VAR, CV and RCU) for the seven variables in wheat cultivars.

The RCU, (*CSS*/*UCSS*)×100, is an index without unit which expressed in terms of percentage and shows what proportion of the sum of squares of the data values is arising out of the squared deviations of them from their mean. In simple terms, what percentage of the difference of data from their mean is explained by the difference of data from each other? Similar to VAR, in the RCU, two numeric values are considered quite similar when their difference is zero and when considered quite different that are exactly symmetric to each other. The values of RCU are ranged between zero and 100 %, when RCU is zero shows there is no variation and all data values are identical to the mean and hence to each other. With the more deviation from the mean and hence to each other, the RCU will be closer to 100 %. In situations the data values are exactly symmetric the RCU reaches to 100 %, therefore the mean and of curse sum total of data is zero. It means that total sum of squares of data is resulting from the difference of them from the mean. Another advantage against both of VAR and CV is the RCU dose not directly affect by sample size, N, particularly repetition of the similar data. Having the minimum and maximum, converts RCU to an absolute measure of variation which represents the level of dispersion *per se*.

As the graphical comparison of the values for VAR, CV and RCU indicates, in terms of the extent of variation is expressed by a criterion, both CV and RCU criteria were similar, while the VAR was not alike with any of these two (Fig. 1). Finding the relationship between the values of RCU and CV will be of great help to understand the actual extent of variation that can be expressed by the RCU. Since CV and RCU are made by using components ∑X, N and ∑X^2^, the relationship between the values of two criteria can be easily concluded. It can be proved that if the CV is equal to 100 % the RCU will be 50 %. The CV=100% shows a sample in which average deviations of data from the mean is equal to the mean, and this amount of dispersion is experientially very high, so that in such cases, the variation is said to be complete. Similarly, if the CV is 50 % (medium variation), then the RCU will be equal to 20 %. And if the CV is 20 % (low variation), then the RCU will be approximately 4 %. Where all data are similar, and there is absolutely no variation, both CV and RCU will be zero. As an experiential rule, therefore, the relationship between the value of RCU and actual variation in the data values is accepted as follows:

| RCU (%) | Variation |
| --- | --- |
| 0 | No |
| Less than 4 | Low |
| 4 to 20 | Medium |
| More than 20 | High |
| 100 | Complete |

The above classification is the special feature of RCU, because allow researchers to quantify and interpret variation just by the RCU rather than comparison other variables RCUs.

The relationship between RCU and CV is well shown for 1000 data sets each consisting of 50 numeric values with mean 500 and different variation in Fig. 2. The figure shows how for CVs more than 400 %, the curve is asymptotic to the unit i.e. RCU=100 %. Now it is easy to qualify variation of the studied variables using RCU. According to the RCU values in Table 2, variation of variable spikes and yield was medium and for other variables was low. There was no variable with high variation.

**Fig 2.**
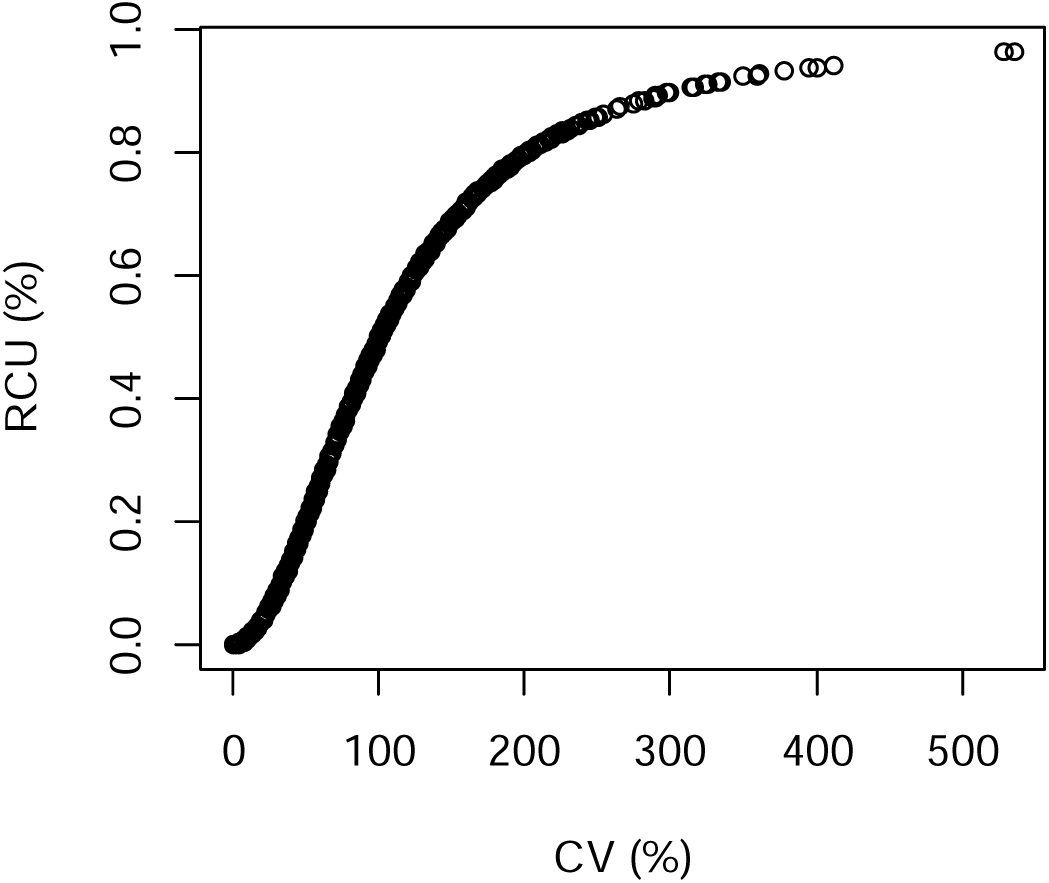
The relationship between RCU and CV for hypothetical data samples with *X*̅ = 500 and different variation.

Generally it can be concluded that, RCU, shows what percentage of a data sum of squares is due to the deviation from the mean, while VAR indicates the average deviations of data from their mean. In comparison to prevalent variation statistics, the RCU has two major advantages; first it is an actual indication of the amount of variation in the sample or population, and the second, using RCU, the variation of two groups of data will be compared easily, if the variation of the first group is higher, its RCU would be more. If there is no difference between the values, RCU will be zero and whatever the difference in data set is more, the RCU will be closer to 100%. When RCU will reach to 100 % that the data are symmetric and the mean (or the sum total of data values) is zero. Also, RCU less than 4, between 4 and 20 and more than 20 % can be attributed to the sets with low, medium and high variation, respectively.

Plant breeders have been realized that diverse plant genetic resources are priceless assets for humankind due to increasingly requirement for feeding a burgeoning world population in future. Presence of diverse plant genetic resources is essential for further improvement of crops by adding more options for the breeders to develop new cultivars. Choice of genetic materials is the most important decision in the plant breeders takes. Breeders choose not only populations with high phenotypic means but also populations having large and useful genetic variation. Therefore genetic variation is important factor to consider when choosing genetic materials to be used as source of cultivars. Some statistics are available to quantify genetic variation that increased the efficiency of germplasm curators and, plant breeders to speed up the crop improvement. Therefore, we believe that RCU provides a simple, accurate and efficient quantity of variation that could be considered a useful criterion beside common statistics like variance.

